# Emergence of oscillations via spike timing dependent plasticity

**DOI:** 10.1101/269712

**Authors:** Sarit Soloduchin, Maoz Shamir

## Abstract

Neuronal oscillatory activity has been reported in relation to a wide range of cognitive processes. In certain cases changes in oscillatory activity has been associated with pathological states. Although the specific role of these oscillations has yet to be determined, it is clear that neuronal oscillations are abundant in the central nervous system. These observations raise the question of the origin of these oscillations; and specifically whether the mechanisms responsible for the generation and stabilization of these oscillations are genetically hard-wired or whether they can be acquired via a learning process.

Here we focus on spike timing dependent plasticity (STDP) to investigate whether oscillatory activity can emerge in a neuronal network via an unsupervised learning process of STDP dynamics, and if so, what features of the STDP learning rule govern and stabilize the resultant oscillatory activity?

Here, the STDP dynamics of the effective coupling between two competing neuronal populations with reciprocal inhibitory connections was analyzed using the phase-diagram of the system that depicts the possible dynamical states of the network as a function of the effective inhibitory couplings. This phase diagram yields a rich repertoire of possible dynamical behaviors including regions of different fixed point solutions, bi-stability and a region in which the system exhibits oscillatory activity. STDP introduces dynamics for the inhibitory couplings themselves and hence induces a flow in the phase diagram. We investigate the conditions for the flow to converge to an oscillatory state of the neuronal network and then characterize how the features of the STDP rule govern and stabilize these oscillations.

## Introduction

Synaptic plasticity is the basis for learning and memory. According to Hebb’s rule [1], which constitutes the foundation for current views on learning and memory, the interaction strength increases between two neurons that are co-activated. When extended to the temporal domain by taking into account the effect of the causal relationship between pre-and post-synaptic firing on the potentiation and depression of the synapse, this rule is known as spike-timing dependent plasticity (STDP). STDP has been identified in various systems in the brain, and a rich repertoire of causal relations has been described [2–12].

STDP can be thought of as a process of unsupervised learning (but see also e.g. [13]). Considerable theoretical efforts have been devoted to investigating the possible computational implications of STDP [14–32]. It was shown that certain STDP rules can give rise to the emergence of response selectivity at the level of the post-synaptic neuron [15, 16, 23], whereas other STDP rules can provide a homeostatic mechanism that balances the excitatory and inhibitory inputs to the cell [26, 31, 33]. For example, in the visual system, modeling studies have shown how spatial correlations together with STDP can develop response selectivity in the form of ocular dominance and directional selectivity [18, 34–39].

The overwhelming majority of computational studies of STDP have focused on the learning dynamics of feed-forward synapses, partly due to the mathematical difficulties associated with investigating learning dynamics in recurrently connected networks. Researchers have only recently been able to address this issue and provide a basic framework for studying STDP in recurrent networks, see e.g. [21–25, 40–42]. A linear approximation is generally used to estimate neuronal response covariance, which serves as the driving force for the STDP dynamics. As a result, the basic non-linear mechanism that can account for the rich neuronal dynamical behavior is largely lacking.

Oscillatory activity has been reported and proposed to play an important role in relation to various cognitive processes including the encoding of external stimuli, attention, learning and consolidation of memory [43–46]. Although the functions of these oscillations remains unresolved, it is clear that neuronal oscillations are abundant in the central nervous system. In addition, oscillatory activity may have a strong effect on STDP since oscillations cause neurons to fire repeatedly with a distinct spike timing relationship. Therefore, in context of development, oscillations and repeated spatiotemporal patterns of activity may play an important role in shaping neuronal connectivity maps [47, 48].

The effect and possible computational role of oscillations on STDP has been addressed in several studies [49–57]. However, in all of these studies the oscillatory activity was either an inherent property of the neuron or inherited via feed-forward connections from inputs that were oscillating and had a clear preferred phase. This raises the question of the origin of these oscillations: are the mechanisms for generating these oscillations genetically hard-wired into the system or can they be acquired via a learning process? A recent numerical study simulating a large scale detailed thalamocortical model argued that oscillations may emerge with STDP [58]. However, the principles that underlie the emergence of oscillations with STDP remain unclear. Under what conditions can STDP give rise by itself to the emergence of oscillatory activity?

Moreover, neuronal oscillations have been reported to show robustness to various perturbations [59]. Can STDP provide a homeostatic mechanism for the regulation and maintenance of specific oscillatory behavior? If so what features of the STDP rule determine oscillatory behavior?

Here we address these fundamental questions by studying the STDP dynamics of the effective couplings between two rival populations. Because STDP dynamics is governed by pre-post correlations it is essential to be able to analyze these correlations and in particular understand how they depend on the synaptic weights themselves. Assuming separation of time scales between fast neuronal responses and a slower learning process, we calculate these correlations in the framework of a rate model for the neuronal responses. Below we describe the rate model and analyze its phase diagram in the plane of the synaptic coupling strengths. Next we define the STDP rule we will utilize and develop a mean field Fokker-Planck approximation for the synaptic weights dynamics in the limit of slow learning rate. This learning dynamics induces a flow on the phase diagram. Thus, the plane of effective interactions, [*J*_12_, *J*_21_], which is the phase diagram that depicts the possible solution for the neuronal responses, is also the phase plane for the STDP dynamics. We then investigate which features of the STDP rule determine whether this flow will converge to a state in which neuronal activity oscillates and how these oscillations are governed by this rule. Finally, we summarize our results and discuss possible outcomes and extensions to the simplified model studied here.

## Results

### The rate model

We explored the STDP dynamics of the effective coupling between two neuronal populations with reciprocal inhibition. We modelled the rate dynamics of the populations as:

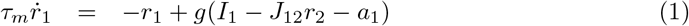

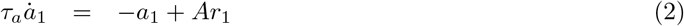

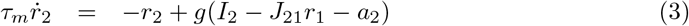

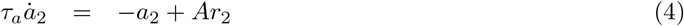

where *r*_*i*_ is the mean rate of population *i* that receives external excitatory input *I*_*i*_. For simplicity we take *I*_1_ = *I*_2_ ≡ *I*. *g*(*x*) is a sigmoidal function and throughout this paper it is taken to be a threshold linear function of its input, *g*(*x*) = ⌊*x*⌋_+_ = *x* for *x* > 0 and 0 otherwise. The terms *a*_1_ and *a*_2_ represent adaptation variables of populations 1 and 2 respectively and parameter *A* denotes the adaptation strength. *J*_*ij*_ ≥ 0 is the strength of inhibition from population *j* to population *i*.

Parameter *τ*_*m*_ is the membrane time constant and *τ*_*a*_ is the adaptation time constant. It is assumed that adaptation is a slower process than the neural response to its input, *τ*_*a*_ > *τ*_*m*_. This model and its variants have been used in the past to model binocular rivalry (Shamir & Sompolinsky unpublished). In the limit of *ϵ* = *τ*_*m*_/*τ*_*a*_ → 0 a complete analytical solution is possible, including the calculation of the limit cycle solution. Unless noted otherwise (mainly in the numerical simulations) the results are given for the *ϵ* → 0 limit. This model and its architecture were chosen for their simplicity and analytical tractability and the fact that they enable oscillatory activity.

### The phase diagram

Fig 1A depicts the phase diagram of the model in the plane of *J*_12_ and *J*_21_ in the limit of *ϵ* → 0. If the inhibition from population 1 to population 2, *J*_21_ is sufficiently strong relative to the adaptation, *J*_21_ > 1 + *A*, there exists a solution that we term *Rival 1*, in which population 1 fully suppresses population 2 (*r*_2_ = 0). Similarly, the *Rival 2* solution, in which population 2 fully suppresses population 1, exists for *J*_12_ > 1 + *A*. The Rival states are stable wherever they exist and may also co-exist.

**Fig 1.**
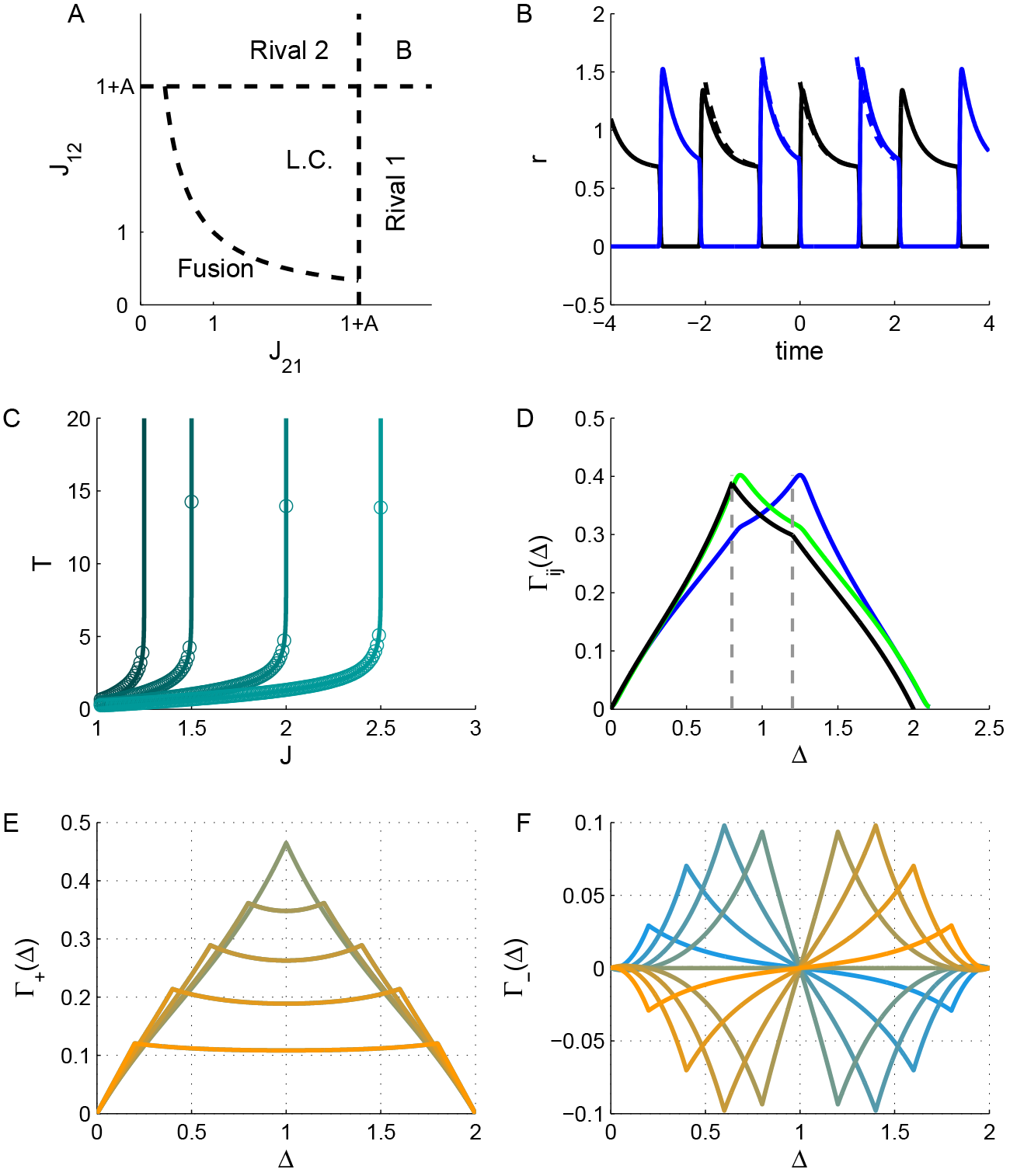
Neuronal dynamics. **A**. The phase diagram. The regions of different types of solutions for the neuronal dynamics are depicted in the (quarter of the) plane of (non-negative) *J*_21_ and *J*_12_. **B**. The limit cycle solution. The firing rate of populations 1 and 2 are plotted in black and blue, respectively, as a function of time (measured in units of *τ*_*a*_) in the anti-phase oscillatory solution with *T*_1_ = 1.2 and *T*_2_ = 0.8, yielding *J*_21_ ≈ 2.36 and *J*_12_ ≈ 1.87 (see Eq (32)). In this specific example we used *I* = 2, *A* = 2, the solid lines show the solution for *ϵ* = 0.01 and the dashed depict the solution in the limit of *ϵ* → 0. **C**. The oscillation period along the diagonal. The oscillation period on the diagonal is shown as a function of the reciprocal inhibition strength for different values of the adaptation strength, *A* = 0.25, 0.5, 1, 1.5 from left to right. Solid lines show the analytical relation of Eq (33) in the *ϵ* → 0 limit. The circles depict the *ϵ* = 0.01 case. **D**. The cross-correlation function. The neuronal cross-correlations Γ_12_ (green and black) and Γ_21_ (blue) are plotted as function of the time difference, Δ (measured in units of the adaptation time constant *τ*_*a*_). The black line depicts the correlations in the *ϵ* → 0 limit, whereas the green and blue lines show the *ϵ* = 0.01 case. Parameters were identical to B. For the *ϵ* = 0.01 case the correlations were evaluated from the numerical solution for the dynamics. **E**. The ‘mean cross-correlation’ function. The mean correlation, Γ_+_, in the limit of *ϵ* → 0, (see Methods) is plotted as a function of Δ for *T* = 2 and different values of the *T*_1_ = *T*[0.1, 0.2,…0.9] shown by color. Note that the plots for *T*_1_ = *x* and *T*_1_ = *T* − *x* overlap. **F**. The ‘difference cross-correlation’. The difference in the cross-correlation, Γ_−_, in the limit of *ϵ* → 0, is plotted as a function of Δ for *T* = 2 and different values of the *T*_1_ = *T* × {0.1, 0.2,…0.9} shown by color from yellow (*T*_1_ = 0.1*T*) to blue (*T*_1_ = 0.9*T*). In E and F *A* = 2 and *I* =2 were used.

For weak reciprocal inhibition, *J*_21_ < 1 + *A* and *J*_12_ < 1 + *A*, there exist a solution in which both populations are active that we term the *Fusion* state. However, this fusion state loses its stability if the inhibition is sufficiently strong, 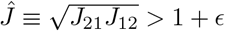. Consequently, there is a region in the phase diagram in which there is no stable fixed point solution. In this region the system relaxes to a limit cycle of anti-phase oscillation, Fig 1B. In the limit of slow adaptation, *ϵ* → 0, one can derive a complete solution for the limit cycle, see Methods. In this case the limit cycle solution has two phases. During the first phase population 1 is dominant and active, *r*_1_ > 0, whereas population 2 is quiescent, *r*_2_ = 0. During the second phase population 2 is dominant and population 1 is quiescent. We denote by *T*_1_ the dominance time of population *i*, and by *T* = *T*_1_ + *T*_2_ the period of the oscillations, see Fig 1B. Along the diagonal of the phase diagram, *J*_12_ = *J*_21_ the dominance times are equal, *T*_1_ = *T*_2_ = *T*/2, and the oscillation period monotonically increases from zero on the boundary of the stable *Fusion* solution, *Ĵ* = 1, to infinity on the boundary of the Rival solutions, *Ĵ* = 1 + *A*, Fig 1C. The dominance time of population 1, *T*_1_, diverges to infinity on the boundary of *Rival 1* state, l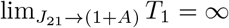, and similarly *T*_2_, diverges on the boundary of *Rival 2* state.

### The correlation function

A central factor that shapes STDP dynamics is the pre-post correlation function. To this end we modelled the spiking activity of neurons in population *i* as independent inhomogeneous Poisson processes with instantaneous rate *r*_*i*_(*t*). Let us denote by *ρ*_*x,i*_(*t*) the spiking activity of neuron *x* in population *i* ∈ {1,2}, which is a Dirac comb of the sum of delta functions at the spike times of the neuron. Thus, the full correlation of different neurons is given by the product of the mean firing rates 〈*ρ*_*x*,1_(*t*)*ρ*_*y*,2_(*t*′)〉 = *r*_1_(*t*)*r*_2_(*t*′). Due to the separation of time scales in the limit of slow learning (see below) the STDP dynamics are driven by the temporal average of the cross correlations. For a periodic solution we define

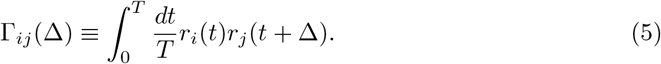

Fig 1D shows the temporal average cross correlation, Γ_*ij*_(Δ), in the asymmetric case, *J*_12_ ≠ *J*_21_, for finite *ϵ* (green and blue) and in the limit of *ϵ* → 0 in black. Note that the main difference is the slight deviation in the oscillation period due to finite *ϵ*, which is more significant at low *T*. A detailed derivation of the cross correlation functions appears in Methods. To analyze the STDP dynamics it is convenient to use the following quantities: Γ_+_(Δ) = (Γ_21_(Δ) + Γ_12_(Δ))/2 and Γ_−_(Δ) = Γ_21_(Δ) − Γ_12_(Δ), as shown in Figs 1E & F, respectively, as a function of the time difference, Δ, for *T* = 2 and different values of *T*_1_ (differentiated by color). In general, Γ_±_(Δ) are periodic functions of time with a period of *T*. Γ_+_(Δ) is an even function of time that is symmetric with respect to *T*/2, whereas Γ_−_(Δ) is an odd function of time that is anti-symmetric with respect to *T*/2. Importantly, on the diagonal of the phase diagram, *J*_12_ = *J*_21_, one obtains that Γ_−_(Δ) = 0.

### The STDP rule

The above analysis was carried out for fixed values for the synaptic weights, assuming that the time scales in which the synaptic weights change are longer than the characteristic time of the neuronal population dynamics, *τ*_*a*_. Next we consider the effect of STDP. We assume that initially synaptic weights are relatively weak (i.e., near the origin of the phase diagram in the *Fusion* state) and examine how activity dependent plasticity shapes its evolution, which induces a flow on the phase diagram. Consequently, the phase diagram of the neuronal activity becomes the phase plane of the synaptic weights. Following Luz and Shamir (2014) the STDP rule is written as the sum of two processes, potentiation and depression,

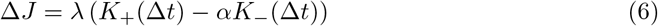

Where Δ*J* is the synaptic weight difference due to pre and post spikes with a time difference of Δ*t* = *t*_post_ − *t*_pre_. The functions *K*_±_(*t*) are the temporal kernels for the potentiation (+) and depression (-) of the STDP rule, respectively, and *α* is the relative strength of the depression. Parameter λ is the learning rate. We assume that the learning process occurs on a slower time scale than the adaptation. Specifically, here we focus on the family of temporally a-symmetric exponential learning rules:

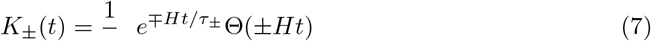

Where Θ(*x*) is the Heaviside step function, and *τ*_±_ denote the characteristic time scales of the LTP and LTD branches of the rule, respectively. The parameter *H* = ±1 governs the nature of the learning rule, with *H* = 1 for a “Hebbian” rule (i.e., potentiating at the causal branch, when the post fires after pre, Δ*t* > 0), and *H* = − 1 for the “Anti-Hebbian” STDP rule. Below we analyze the mean field approximation in the limit of λ → 0.

### The mean field Fokker-Planck dynamics

Changes to the synaptic weights due to the plasticity rule of equation (6) in short time intervals occur as a result of either a pre or post-synaptic spike during this interval. Thus, we obtain

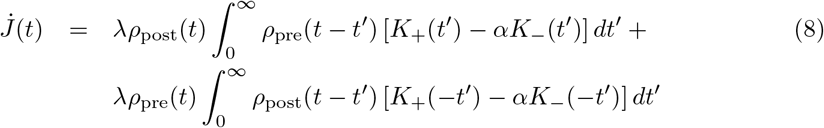

The mean-field approximation is obtained in the limit of λ → 0, where the right hand side of equation (8) can be replaced by its temporal mean due to the averaging of the slow learning dynamics, yielding

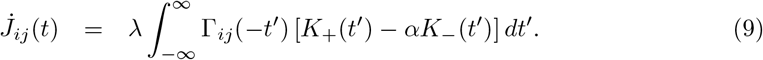

In regions of the phase diagram where a stable fixed point solution exists, i.e., 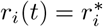, the correlation function, Γ, is given by the product of the time independent means, 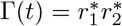, and one obtains that 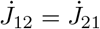. As the firing rates are non-negative and the temporal kernels of the potentiation and depression, *K*_±_, have an integral of one, the sign of 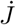 is determined by 1 − *α*. As a corollary, the synaptic weights will flow towards the region of limit cycle solution from initial conditions close to the origin in the phase diagram if *α* < 1. This result holds for any choice of temporal structure for the STDP rule. Note that a similar condition (*α* < 1) was assumed for inhibitory plasticity in Luz and Shamir (2012). Thus, initial conditions of weak synaptic coefficients (*J*_*ij*_ close to the origin) will flow towards the region of the limit cycle solution and will enter it near the diagonal, *J*_21_ = *J*_12_.

In the region of the limit cycle the STDP dynamics do not necessarily flow in parallel to the identity line, but rather depend on the specific limit cycle solution and on the temporal structure of the STDP rule. It is convenient to formulate the STDP dynamics in terms of *J*_+_ ≡ (*J*_21_ + *J*_12_)/2 and *J*_−_ ≡ *J*_21_ − *J*_12_, yielding

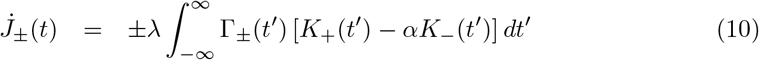

On the diagonal, *J*_12_ = *J*_21_, due to the symmetry of the limit cycle solution Γ_12_(*t*) = Γ_21_(*t*) and as a result 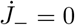. The mean correlation, Γ_+_, on the other hand, is a positive even function of time with a period of *T*. Near the boundary of stable *Fusion*, the oscillation frequency diverges, *T* → 0. In this limit (for *ϵ* → 0) the limit cycle solution for the neuronal responses will approach a square wave solution (with 50% duty cycle on *J*_12_ = *J*_21_) transitioning between 0 and 2*I*/(2 + *A*) in anti-phase. The mean correlation function, Γ_+_(Δ), will approach a triangular wave starting at 0 for Δ = 0 and peaking at 2*I*^2^/(2 + *A*)^2^ for Δ = *T*/2. Consequently, for *T* → 0, the integral on the right hand side of equation 10 will be dominated by the DC component of Γ_+_, yielding 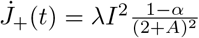 in this limit. Hence, the same condition that allows the STDP dynamics to enter the limit cycle region from the *Fusion* region will also cause it flow in the *J*_+_ direction after entering the Limit cycle region.

Eqs (10) provide two non-linear equations for *J*_+_ and *J*_−_ that are coupled in a non trivial manner via the dependence of the correlations on the synaptic weights. However, on the diagonal of the phase diagram the situation is simplified; since 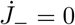 the problem is reduced to a one dimensional flow. To analyze the dynamics of *J*_+_ on the diagonal it is convenient to write it as the sum of two terms:

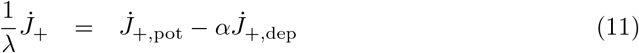

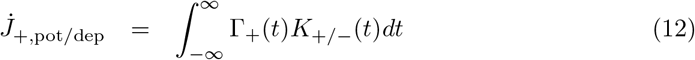

Figs 2A & B show 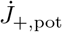 and 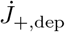, respectively, on the diagonal as a function of the oscillation period, *T* (note that *T* is a function of *Ĵ*, see Fig 1C), for different values of *A* (differentiated by color). As can be seen from the figure, 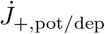 decreases monotonically from the value of *I*^2^/(2 + *A*)^2^ at *T* = 0 to 0 as *T* → ∞ at *J*_12_ = *J*_21_ = 1 + *A* (at the crossing to the bi-stable region). Due to the symmetry of the mean cross-correlation function, Γ_+_(*t*) = Γ_+_(*−t*), one obtains that 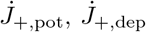 and 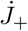 are independent of *H*. Thus the results of Fig 2 hold for both Hebbian and anti-Hebbian plasticity rules. Moreover, 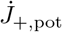 and 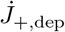 only differ by the time constant of *K*_±_. Fig 2C shows 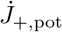 as a function of the oscillation period, *T*, for different values of *τ*_+_ (depicted in color). All the curves decrease monotonically to zero, albeit with a different time scale; consequently, if *τ*_+_ < *τ*_−_ then 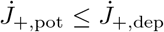 and there is equality only at *T* = 0 (on the boundary of stable *Fusion*).

**Fig 2.**
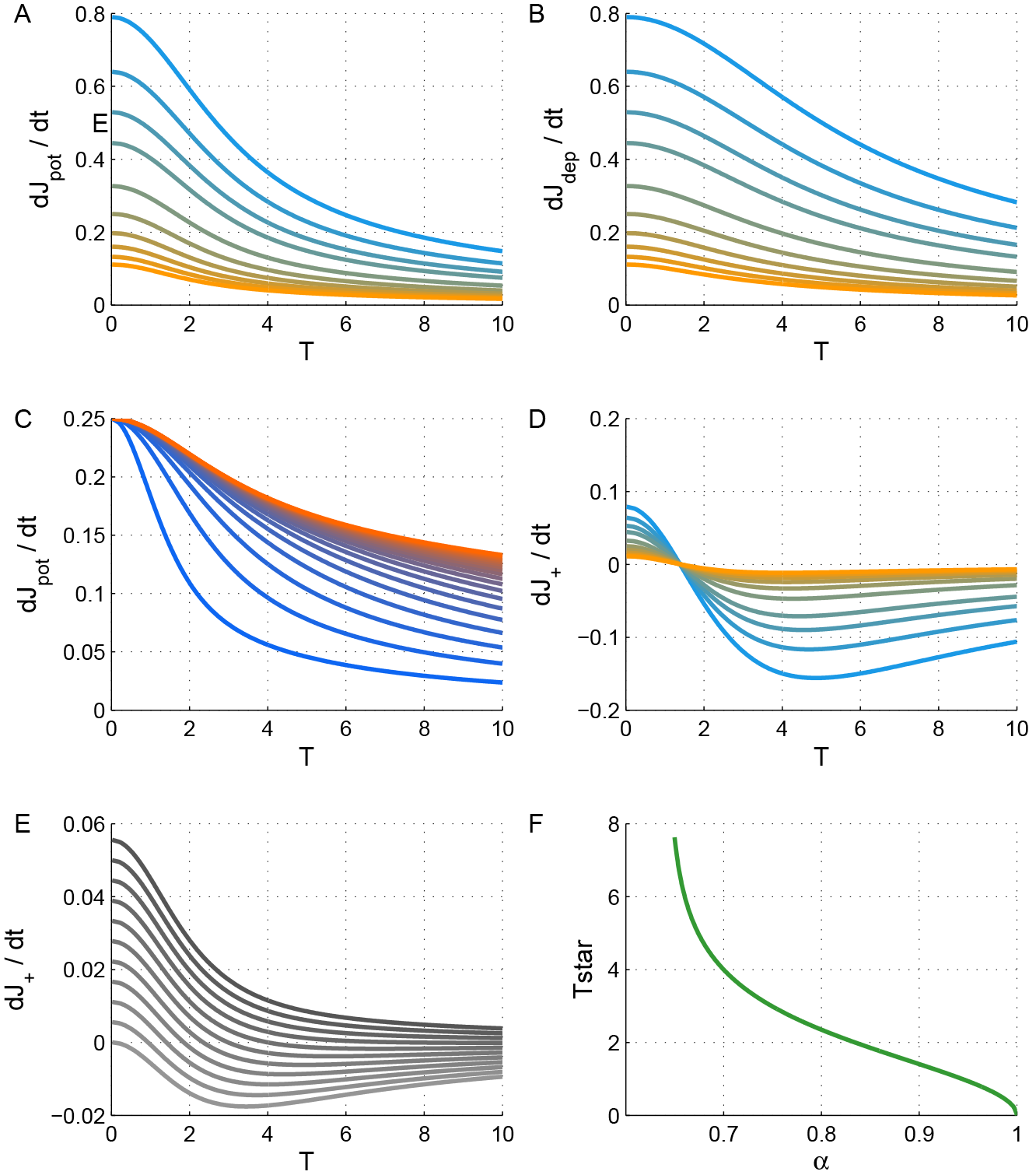
The dynamics of *J*_+_ along the diagonal. **A**. potentiation term, 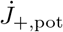, of the mean synaptic weights, *J*_+_, equation (12), is shown as a function of the oscillation period along the diagonal for different values of *A* = 1/4, 1/2, 3/4, 1, 3/2,…4 (from top at low *A* values to bottom). **B**. The depression term, 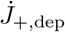, of the mean synaptic weights, *J*_+_, Eq (12), is shown as a function of the oscillation period along the diagonal for different values of the adaptation strength, *A* (as in A). **C**. The effect of the STDP time constant. The potentiation term, 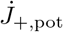, is shown as a function of the oscillation period along the diagonal for different values of *τ*_+_ = 1/4, 1/5,…5, by different colors from blue (low *τ*_+_) to red. Here *A* = 2 was used. **D** The *J*_+_ dynamics along the diagonal. The value of 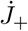 is shown as a function of the oscillation period along the diagonal for different values of *A* using the same values and color code as in A, using *α* = 0.9. **E**. The effect of the relative strength of depression. The value of 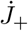 is plotted as a function of the oscillation period along the diagonal for different values of *α* = 0.5, 0.55,…1 from top (*α* = 0.5) to bottom (with *A* = 4). **F**. Oscillation period at the STDP fixed point. The ‘learned’ oscillation period, *T*\*, is shown as a function of *α*. In all panels *I* = 2 was used, and λ = 1 was taken in D and E - for purpose of illustration. Unless otherwise stated (C) were *τ*_+_ = 0.5 and *τ*_−_ = 1 used. All units of time were measured in units of *τ*_*a*_.

The dynamics of *J*_+_ along the diagonal are determined by the weighted sum of both 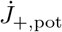 and 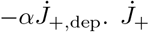 will be positive for *α* < 1 for small *T* - near the crossing from the *Fusion* region. For *τ*_+_ < *τ*_−_ and 1 > *α* > *α*_*c*_(*τ*_+_, *τ*_−_) (see Methods), 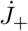 will change its sign at *T*\*; thus, the fixed point (note 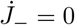 on the diagonal) at *T*\* will be stable along the *J*_+_ direction. This scenario is illustrated in Fig 2D that shows 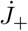 on the diagonal as a function of *T* (for different values of *A*, depicted by color). Interestingly, for this choice of exponential kernels for the STDP rule, the fixed point does not depend on the adaptation strength, *A*. The oscillation period at the fixed point, *T*\*, is zero for *α* = 1 and diverges as *α* approaches a critical value *α*_*c*_(*τ*_+_, *τ*_−_), Fig 2E & F. For fixed *α* ≤ 1 and *τ*_+_, *T*\* is minimal for *τ*_−_ → ∞, increases monotonically as *τ*_−_ decreases and will diverge for a critical value *τ*_−,c_ < *τ*_+_ such that *α*_*c*_(*τ*_+_, *τ*_−_) = *α*. For *τ*_−_ < *τ*_−,*c*_ (and *α* ≥ 1) there will be no fixed point along the diagonal and the STDP dynamics along the diagonal will flow outside of them limit cycle region.

The stability of the STDP fixed point requires stability in the *J*_−_ direction as well. On the diagonal 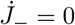. A small perturbation in the direction of *J*_−_ will affect *J*_−_ dynamics via the cross correlation term Γ_−_(Δ), Eq (10). The cross correlations depend on the synaptic weight via the dominance times, *T*_1_ and *T*_2_. Hence, for a small perturbation around the diagonal, Δ*J*_−_ = *J*_−_, one obtains

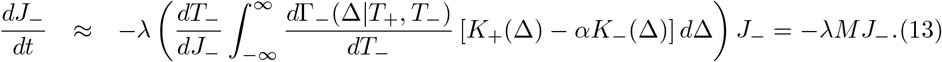

The geometry of the phase diagram (Fig 1A) reveals that increasing (decreasing) *J*_−_ results in advancing towards the Rival 1 (Rival 2) region, and consequently increasing *T*_1_(*T*_2_) and (decreasing) *T*_−_; hence, 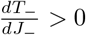. As above, we can define 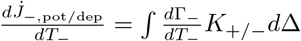. Figs 3A & B show 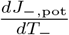 and 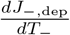 respectively, along the diagonal as a function of *T* for different values of *A* (depicted by color) for Hebbian STDP, *H* − 1. In contrast with Γ_+_(Δ) that was always positive and an even function of Δ, Γ_−_(Δ) and similarly 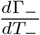 is not necessarily positive and an odd function of Δ. Consequently, 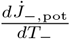 and 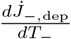 in Fig 3A & B have different signs. The value of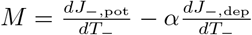 is depicted along the diagonal as function of the oscillation period, *T*, for different values of *A* (differentiated by color) and *α* (shown by grey level) in Fig 3C &D, respectively. Here, *M* is positive, and as a result, STDP dynamics will be stable with respect to fluctuations in the *J*_−_ direction for Hebbian plasticity.

**Fig 3.**
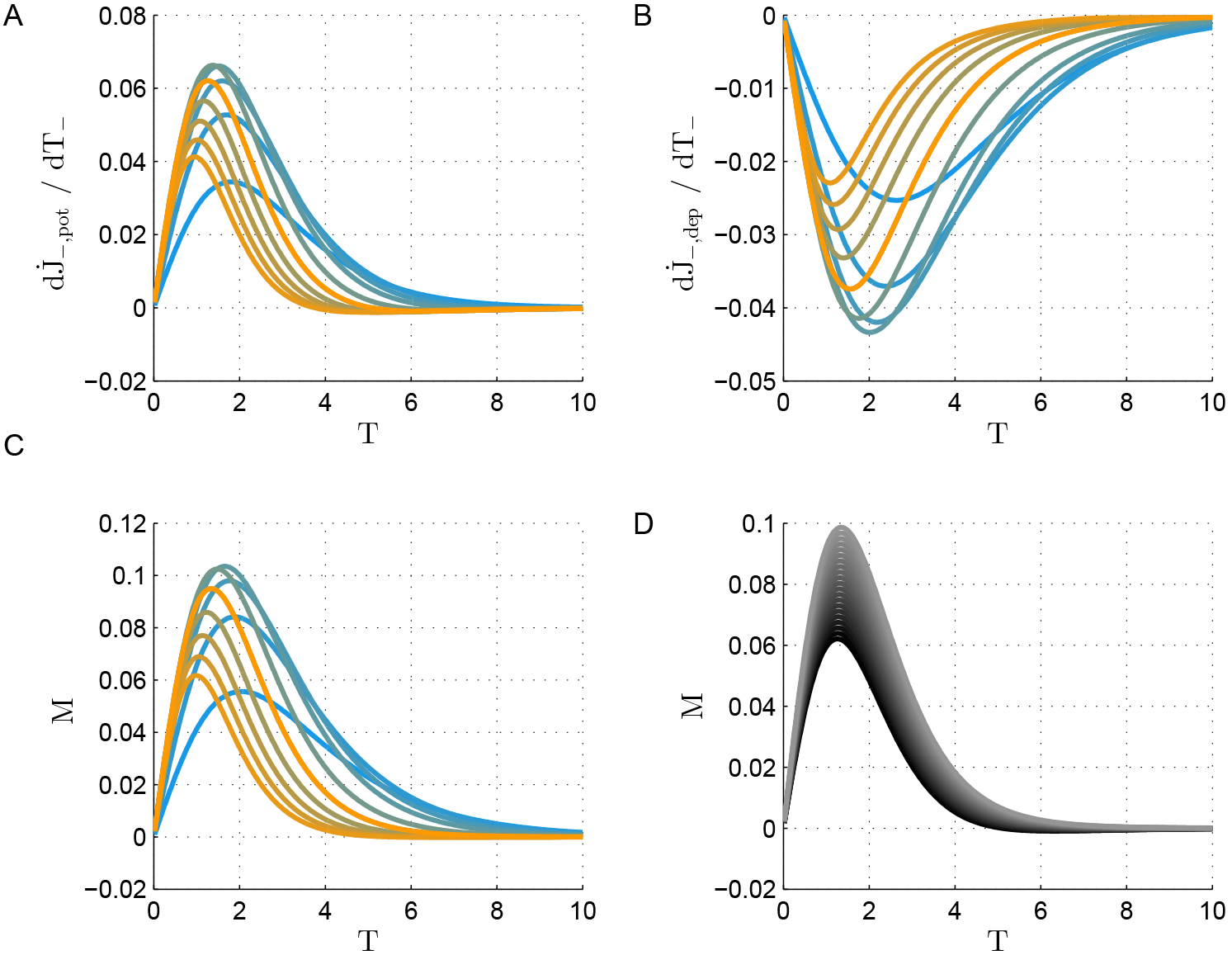
Stability in the*J*_−_direction along the diagonal. **A**. The value of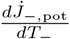 is shown as a function of *T* along the diagonal of the phase-diagram in the *Limit cycle* region for different values of *A* = 1/4, 1/2, 3/4, 1, 3/2,…4 (from top at low *A* values to bottom). Here *τ*_+_ = 0.5 was used. All units of time were measured in units of *τ*_*a*_. **B**. The value of 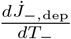 is shown as a function of *T* for different values of the adaptation strength, *A* (as in A). Here *τ*_−_ = 1 was used. **C** *J*_−_ dynamics along the diagonal. The value of 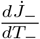 is shown as a function of the oscillation period along the diagonal for different values of *A* (same values and color code as in A), using *α* = 0.9. **D**. The effect of the relative strength of depression. The value of 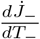 is plotted as a function of the oscillation period along the diagonal for different values of *α* = 0.5, 0.55,…1 from bottom (dark, *α* = 0) to top (bright, *α* = 1), using *A* = 2.

In contrast with Γ_+_, Γ_−_ is an odd function of time, Γ_−_(*t*) = −Γ_−_(*t*). Switching from the Hebbian plasticity rule, *H* = 1, to anti-Hebbian, *H* = −1, will change the sign of 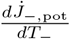, 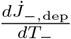 and of *M*. As a result a fixed point (on the diagonal) that is stable in the *J*_−_ direction for Hebbian plasticity will be unstable for anti-Hebbian plasticity and vice versa. Fig 4 shows the flow induced by the STDP on the phase diagram for the (A) Hebbian and (B) Anti-Hebbian learning rules. As can be seen, the Hebbian learning rule is unable to converge to a state that allows oscillatory activity. In contrast, the Anti-Hebbian STDP generate symmetric (*T*_1_ = *T*_2_) anti-phase oscillatory activity, in which the oscillation period is determined and controlled by the relative strength of depression, *α*. This specific learning rule provides robustness with respect to the strength of adaptation, *A*. Fluctuations in *A* do not affect the period of the oscillation.

**Fig 4.**
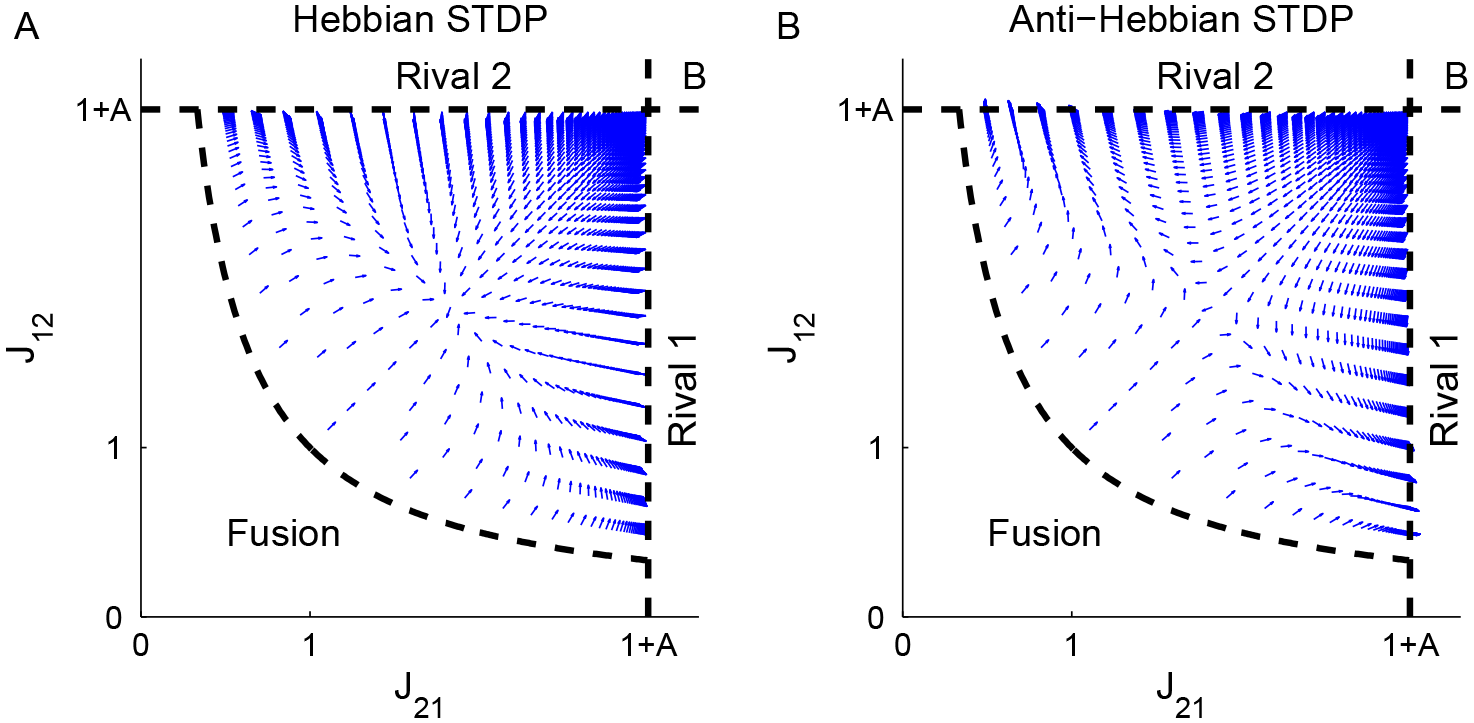
The flow on the phase diagram. The direction of the dynamic flow, i.e., the normalized vector 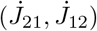, is shown in the *Limit cycle* region of the phase diagram for **A**. Hebbian plasticity, *H* = 1 in Eq (7), and **B**. Anti-Hebbian plasticity, *H* = −1. The parameters used here were: *A* = 2, *τ*_+_ = 0.5 and *τ*_−_ = 1

## Discussion

We examined whether oscillatory activity can emerge via an unsupervised learning process of STDP. Our main result is that under a wide range of parameters, oscillatory activity can develop via STDP. Specifically, we found that to develop the capacity for oscillatory activity the STDP rule must obey the following conditions (*i*) a bias towards potentiation, *α* < 1 will lead the system into the oscillatory region of the phase diagram, (*ii*) a longer characteristic time for depression than for potentiation, *τ*_−_ > *τ*_+_, will enable the existence of a fixed point on the diagonal that can be governed by the exact value of *alpha*, and (*iii*) the stability of the fixed point in the orthogonal direction is governed by the ‘Hebbianity’ of the plasticity rule. STDP may also provide a mechanism for selecting and stabilizing the desired oscillations; for example, oscillation frequency can be governed and manipulated by the relative strength of the depression, *α*, or changes in the time constants of the STDP rule, *τ*_±_. Disruption of the STDP rule may result in changes to the oscillation frequency.

Analysis of STDP dynamics in recurrent networks is challenging. To facilitate the analysis we used the framework of a simplified model for the neuronal responses, and explored the learning dynamics of the *effective* couplings between the two populations. We assumed a separation of three time scales *τ*_*m*_ ≪ *τ*_*a*_ ≪ λ ^−1^. The separation of the neuronal time constant from that of the adaptation enabled us to obtain an analytic expression for the temporal correlations that drive the STDP dynamics. The assumption that long term synaptic plasticity occurs on a longer time-scale allowed us to consider STDP dynamics as a flow on the phase diagram.

The interplay of short and long term plasticity processes deserves consideration. Oscillations would not be possible in this model without the short term plasticity; here, adaptation. Thus, short term plasticity has a major role in shaping the temporal structure of the neuronal cross-correlations, Γ_*ij*_(*t*) that drive the STDP dynamics, which in turn, may or may not converge to a state that allows this oscillatory behaviour.

The reflection of the flow on the phase diagram with respect to the diagonal when reflecting the STDP rule with respect to time stems from the inherent symmetry of the cross correlation function which drives the dynamics (Γ_*ij*_(Δ) = Γ_*ji*_(−Δ)); hence, it is general and holds regardless of the choice of model. Certain other assumptions can be easily relaxed. For example, we assumed symmetry between the two competing populations. However, using the (threshold) linearity of our model one can easily rescale the neuronal responses to allow for different inputs and adaptation strengths. On the other hand, the independence of the fixed point, *T*\*, on the adaptation strength, *A*, is specific to this model and for the choice of an exponentially decaying STDP rule.

A central assumption in this study was the choice of (a reciprocal inhibition) architecture. This choice was made to obtain a model that can be fully analyzed. However, the choice of architecture (including the short-term-plasticity mechanism) shapes the phase diagram, allows for the different regions of dynamical solutions (fixed points, In/Out of/Anti -phase oscillations, etc.) and determines the cross correlations. Consequently, the effect of the network architecture on STDP dynamics should not be underestimated. Because this effect is highly non-linear, one cannot generalize these results to other architectures in a straightforward manner. Nevertheless, the approach delineated here, namely, studying the induced flow on the phase diagram of the system, can be applied to other models in the limit of slow learning rate.

## Methods

### Phase diagram and limit cycle calculations

#### The fixed points of the dynamics

We distinguish two types of fixed points: *Rival* states, in which one population fully suppresses the other, and *Fusion*, in which both populations are active.

**The Rival states.** The *Rival-1* solution assumes 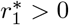 and 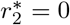, yielding 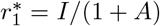, 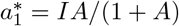 and 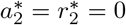. The existence condition for this solution is that the net input to population 2, *I* − *J*_21_*r*_1_ − *a*_2_ is non-positive, at the fixed point, *J*_21_ ≥ 1 + *A*. This solution is always stable where it exists.

**The Fusion state.** The *Fusion* solution assumes 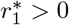 and 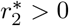, yielding

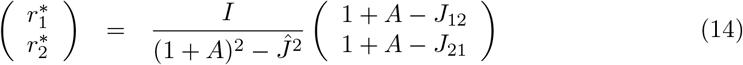

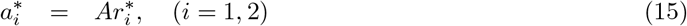

where 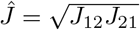. The existence of the *Fusion* solution requires the inputs of both populations to be non-negative. For *Ĵ*^2^ < (1 − *A*)^2^ the existence condition requires *J*_12_ ≤ 1 + *A* and *J*_21_ ≤ 1 + *A*(bottom left square in the phase diagram, Fig 1A, where no *Rival* solution exists). By contrast, for *Ĵ*^2^ > (1 - *A*)^2^ the existence condition requires *J*_12_ ≥ 1 + *A* and *J*_21_ ≥ 1 + *A* (the region in the phase diagram where both *Rival* solutions exist). However, the *Fusion* state is not always stable. By performing standard stability analysis around the *Fusion* fixed point we expand the dynamics around the fixed point to a leading order in the fluctuations

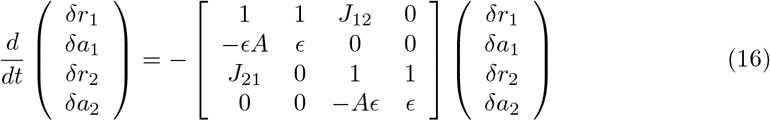

where *δx* ≡ *x* − *x*\*, yielding the four eigenvalues for the stability matrix:

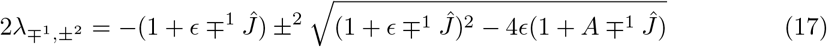

The sum of the pair of eigenvalues λ_+^1^,±^2^_ is − *Ĵ* − (1 + *ϵ*) < 0 and their product is *ϵ*(1 + *A* +^1^ *Ĵ*) > 0; hence, these eigenvalues are always stable. On the other hand, for the pair of eigenvalues λ_−1,±2_ the sum is +*Ĵ* − (1 + *ϵ*), which is negative if and only if inhibition is sufficiently week, *Ĵ* < 1 + *ϵ* (in that case their product will also be positive, assuming *ϵ* is small). Thus, the *Fusion* state looses its stability when reciprocal inhibition becomes sufficiently strong, *Ĵ* > 1 + *ϵ*.

**The Limit Cycle solution.** In the region of the phase diagram where no stable fixed point exists the network dynamics relaxes to anti-phase oscillations. Below we provide a detailed solution for the limit cycle in the limit of *ϵ* → 0. The limit cycle is solved using the anti-phase oscillations ansatz. First the neuronal dynamics is solved for each phase, where the dynamics are linear. This provides a piecewise solution with several parameters to be determined. Then we apply two sets of constraints: periodicity and transition.

Assuming the anti-phase oscillations ansatz we separate the cycle into two phases. During *phase-1* population 1 is dominant and fully suppresses population 2, for times *t* ∈ (0, *T*_1_). In the limit of slow adaptation, *ϵ* → 0, dynamics during *phase-1* are given by:

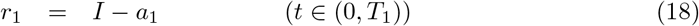

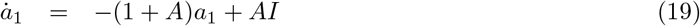

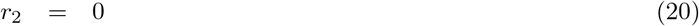

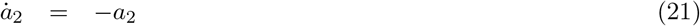

where we measure time in units of *τ*_*a*_. eqs (18)-eqs(21) can be easily solved, yielding

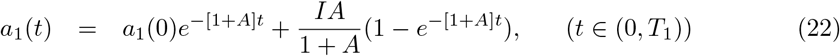

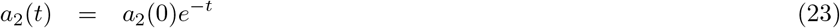

Similarly, during *phase-2*, when population 2 is dominant and fully suppresses population 1, *t* = *t*′ +*T*_1_ ∈ (*T*_1_, *T*_1_ + *T*_2_), we obtain

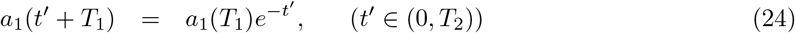

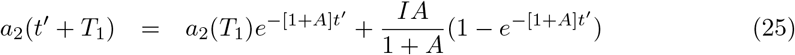

Continuity of the adaptation variables, *a*_*i*_, dictates that, for example, the initial conditions of Eq (25), *a*_2_(*T*_1_), will be given from Eq (23), *a*_2_(*T*_1_) = *a*_2_(0)*e*^−*T*1^. We now need to determine four parameters: *a*_1_(0), *a*_2_(0), *T*_1_ and *T*_2_. These parameters are determined by two sets of constraints. One is periodicity, namely

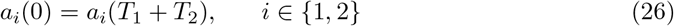

yielding,

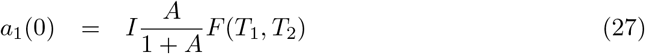

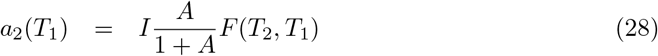

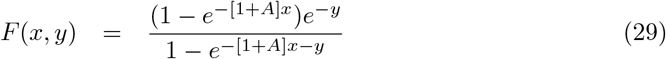

The second set of constraints is given by the transition conditions. Specifically, the transition time from *phase-1* to *phase-2* at *T*_1_ is not arbitrary; rather, *T*_1_ is a special point in time in which population 2 is released from being fully suppressed, such that, the net input to population 2 changes its sign from negative to positive; thus,

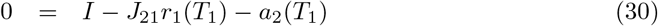

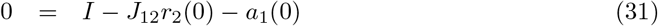

which provides implicit equations for the dominance times, *T*_1_ and *T*_2_,

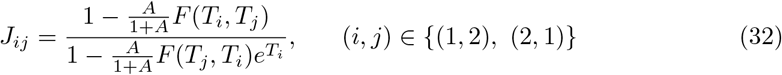

Using Eq (32), and taking the limit of *T*_1_ → ∞, we obtain *J*_21_ → 1 + *A*. Thus, the dominance time of population *i*, *T*_*i*_, diverges on the boundary of *Rival-i*. Taking the limit of *T*_1_, *T*_2_ → 0 such that *T*_1_/*T*_2_ = *β* yields 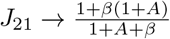 and from symmetry 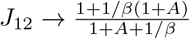, which obeys *J*_12_*J*_21_ → 1; hence, the limit of zero oscillation period is obtained on the boundary of stable *Fusion* (note that these calculations were done for *ϵ* → 0).

On the diagonal, *J*_12_ = *J*_21_ ≡ *Ĵ*, dominance times are equal, *T*_1_ = *T*_2_ = *T*/2,

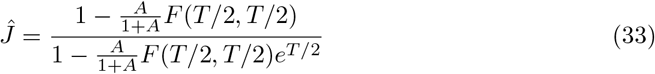

Consequently, the oscillation period, *T*, increases monotonically along the diagonal of the phase-diagram from zero at the transition to *Fusion* (*Ĵ* = 1) to infinity at the transition to the *Rival* states (*Ĵ* = 1 + *A*)

### Calculation of the cross-correlation function

Calculation of the (temporally averaged) cross-correlation function, Eq (5), is done using the analytical solution for the neuronal responses in the limit of slow adaptation, *ϵ* → 0 When the system relaxes to a fixed point solution,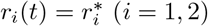, the cross-correlations are constant in time,

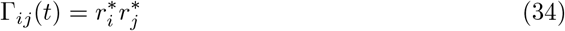

Thus, correlations will be zero in the *Rival* states; hence, there will be no STDP. In the *Fusion* state the cross-correlations will be symmetric, Γ_12_(*t*) = Γ_21_(*t*). As a result, the STDP dynamics for *J*_12_ and *J*_21_ will be identical and the flow will be in the uniform direction, parallel to the diagonal line.

At the *Limit cycle* we use the analytical solution, eqs (22)-(29), to calculate the cross-correlations in a straightforward manner. For Δ ∈ [0, min{*T*_1_, *T*_2_}] we obtain

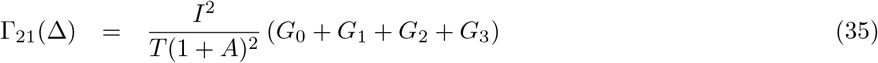

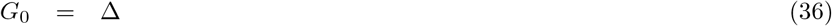

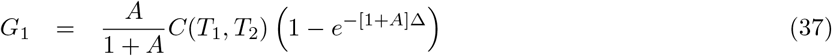

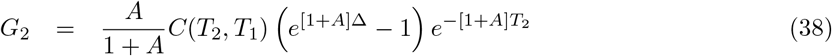

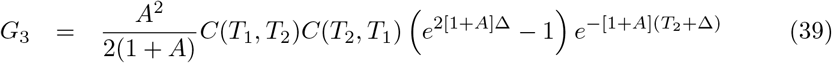

where

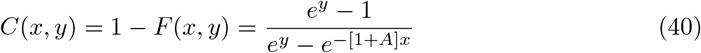

For Δ > min{*T*_1_,*T*_2_} assuming without loss of generality that *T*_1_ ≥ *T*_2_

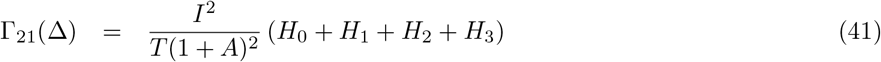

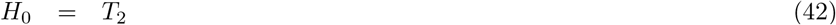

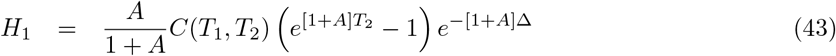

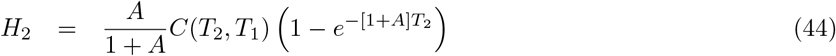

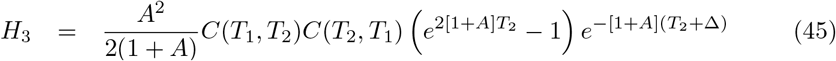

Along the diagonal, on the edge of the stable *Fusion* state region, *T* → 0, the cross-correlation will resemble a triangular chainsaw function (in the *ϵ* → 0 limit) with period *T* and peak 2*I*^2^/(2 + *A*)^2^. Consequently, as *T* goes to zero, the overlap between the cross-correlation function and the STDP rule will be governed by the DC component, yielding

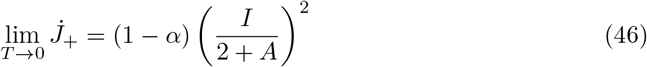

The above expressions for the cross-correlations were given in terms of the dominance times, {*T*_*i*_} instead of the effective couplings *J*_*ij*_. The translation to the synaptic weights from the dominance times is possible by Eq (32). However, because we were interested in studying the ability to learn and stabilize a specific oscillatory activity it was more convenient to think of the dynamics in terms of the dominance times. Similarly, to consider stability with respect to the *J*_−_ direction we utilized the derivative of Γ_−_ = Γ_21_ − Γ_12_ with respect to Δ*T* = *T*_1_ − *T*_2_. On the diagonal, *T*_1_ = *T*_2_ ≡ *T̄*

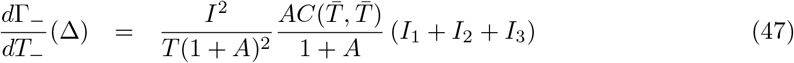

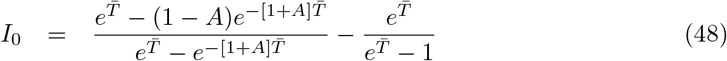

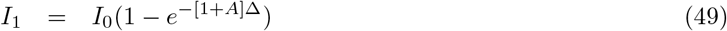

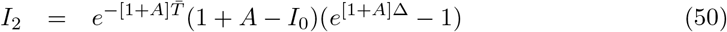

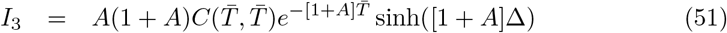

### Calculation of *α*_c_

On the diagonal *T*_1_ = *T*_2_ = *T*/2, in the limit of slow oscillations, *T* → ∞ one obtains

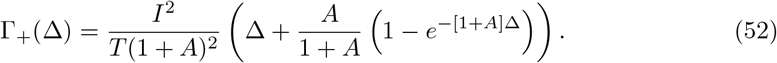

Using Eq (52) yields

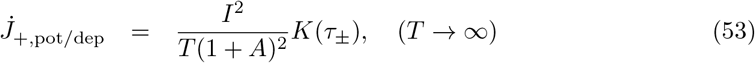

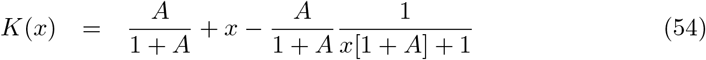

Hence, if *α* is less than a critical value *α*_*c*_ = *K*(*τ*_+_)/*K*(*τ*_−_), 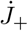 will always be positive (along the diagonal). On the other hand, if *α* is larger than *α*_*c*_ then 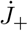 will always be negative for large *T*, and a fixed point will exist if *α* < 1.

## Acknowledgment

The firing rate model of two populations with reciprocal inhibition and slow adaptation has been developed in the past by H. Sompolinsky to model binocular rivalry.

